# On the interactions of the receptor-binding domain of SARS-CoV-1 and SARS-CoV-2 spike proteins with monoclonal antibodies and the receptor ACE2

**DOI:** 10.1101/2020.04.05.026377

**Authors:** Carolina Corrêa Giron, Aatto Laaksonen, Fernando L. Barroso da Silva

**Affiliations:** Universidade Federal do Triângulo Mineiro, Departamento de Saúde Coletiva, Rua Vigário Carlos, 38025-350 – Uberaba – MG, Brazil; Universidade de São Paulo, Departamento de Ciências Biomoleculares, Faculdade de Ciências Farmacêuticas de Ribeirão Preto, Av. café, s/no – campus da USP, BR-14040-903 – Ribeirão Preto – SP, Brazil; Department of Materials and Environmental Chemistry, Arrhenius Laboratory, Stockholm University, SE-106 91 Stockholm, Sweden; State Key Laboratory of Materials-Oriented and Chemical Engineering, Nanjing Tech University, Nanjing, 210009, P. R. China; Centre of Advanced Research in Bionanoconjugates and Biopolymers, Petru Poni Institute of Macromolecular Chemistry Aleea Grigore Ghica-Voda, 41A, 700487 Iasi, Romania; Department of Engineering Sciences and Mathematics, Division of Energy Science, Luleå University of Technology, SE-97187 Luleå, Sweden; Department of Chemical and Biomolecular Engineering, North Carolina State University, Raleigh, North Carolina 27695, United States

## Abstract

A new betacoronavirus named SARS-CoV-2 has emerged as a new threat to global health and economy. A promising target for both diagnosis and therapeutics treatments of the new disease named COVID-19 is the coronavirus (CoV) spike (S) glycoprotein. By constant-pH Monte Carlo simulations and the PROCEEDpKa method, we have mapped the electrostatic epitopes for four monoclonal antibodies and the angiotensin-converting enzyme 2 (ACE2) on both SARS-CoV-1 and the new SARS-CoV-2 S receptor binding domain (RBD) proteins. We also calculated free energy of interactions and shown that the S RBD proteins from both SARS viruses binds to ACE2 with similar affinities. However, the affinity between the S RBD protein from the new SARS-CoV-2 and ACE2 is higher than for any studied antibody previously found complexed with SARS-CoV-1. Based on physical chemical analysis and free energies estimates, we can shed some light on the involved molecular recognition processes, their clinical aspects, the implications for drug developments, and suggest structural modifications on the CR3022 antibody that would improve its binding affinities for SARS-CoV-2 and contribute to address the ongoing international health crisis.

## INTRODUCTION

The SARS-CoV-2, virus recently found in Wuhan, Hubei province, China and officially named by the World Health Organization (WHO),^1^ has already spread through China from all continents (more than 168 countries), with 1,133,758 confirmed cases globally and 62,784 deaths (data as reported by Central European Time 5 April 2020). Due to the pandemic, the disease is not only affecting the health services, but also the economy in a global scale, interfering in the widespread displacement of people, tourism, local and even international markets. Once China’s economy is a worldwide reference, its disruption leads to a global impact in the supply chains and the production itself.^1–3^

The *Coronaviridae* family, to which SARS-CoV-2 belongs, includes a large variability of viruses and became recognized in the spring of 2003, when a human coronavirus caused severe acute respiratory syndrome (SARS).^4–7^ Based on phylogenetic analysis, the SARS-CoV-2 is classified as a lineage B betacoronavirus^8^ and belongs to the same group as SARS-CoV-1 and HKU9-1, the bat coronavirus, demonstrating wide similarity with both genetically^9^ (96,2% of sequence identity with HKU9-1).^10^ The transmission was confirmed to be human-to-human once several medical care personnel and relatives got infected,^8,9^ but it is believed that it all started with an animal host, may it be a bat or another intermediate host.^7,9,11,12^ Even though the virus usually does not cause severe damage to the body, as will be explained below, the major concern is its high infectivity and pathogenicity.^8,12,13^

The COVID-19 disease, caused by SARS-CoV-2 virus,^14^ generally causes mild upper respiratory tract infections, resulting in fever and cough, yet it can also affect the lower respiratory tract.^12,15,16^ SARS-CoV-2, on the other hand, usually remains asymptomatic in an early stage and then manifests itself with dyspnea, severe pneumonia and even death,^17^ with fatality rates of about 10%.^18^ Although many groups of researchers are combining their efforts to solve the mysteries of the new virus, some issues are still uncertain. Examples of these queries are the virus’ incubation period, that may be longer than the 14 days scientists believed it to be previously, and the fatality rates for each age range.^19^

Despite the fact that the virus’ molecular mechanism is partially unknown, the SARS-CoV-2 has proteins, such as the Spike (S) glycoprotein, that densely decorates the viral external surface and can potentially be a key target for the development of vaccines and therapeutic antibodies (Abs).^8,20–22^ Due to the similarity of the receptor binding domain (RBD) in SARS-CoV-2 and SARS-CoV-1, the first strategy that has been used is to search for Abs that succeed interacting with both, once SARS-CoV-1 has been more widely studied. However, preliminary experimental studies have shown that many Abs that successfully interact with SARS-CoV-1 do not bind with SARS-CoV-2.^8^

The spike protein, which is responsible for the “corona” (latin word for crown) appearance in all coronaviruses, is a type I glycoprotein that has an especial role in the interaction between the virus and the host cell. This protein attaches itself to specific cellular receptors and suffers a conformational change that enables the fusion of the virus and the cell.^4,23,24^ Studies have shown that the SARS-CoV-2’s S RBD protein interacts strongly with the Angiotensin-converting enzyme 2 (ACE2).^9,24,25^ Therefore, aiming to develop better diagnosis tools, vaccines and therapeutic Abs, it was measured the competition of mAbs and the ACE2 for the binding to SARS-CoV-2 (named before 2019-nCoV^8,26^) RBD protein in order to enlighten the binding epitopes of these Abs.^8,19^

The focus of this article is to initially reproduce the observations of previous laboratory experiments by a theoretical approach. Secondly, we aim to contribute with the understanding of the molecular mechanisms involved in the SARS viral infection, and finally to show how to apply this knowledge to design new functional molecules. To achieve these goals, it was tested by constant-pH simulation methods the complexation between the S RBD proteins of SARS-CoV-1 and SARS-CoV-2 with the fragments of the monoclonal Abs (mAbs) 80R, CR3022, m396 and F26G29, measuring their binding affinities and quantifying the titratable amino acids that are involved in these interactions. Thus, using a theoretical method recently proposed to identify “electrostatic epitopes”,^27^ it is possible to identify the similarities and differences between these molecular complexes, and to map their origin and possible biological implications.

Another aspect discussed in this research is the interaction between the S RBD protein from these viruses and the ACE2 in order to discover if the S RBD protein binds to either of them with higher affinity, because, if so, an antibody (Ab) might have smaller chances of binding. All this information together provided important insights to design more specific and effective neutralizing Abs which is relevant for the future prevention and treatment of this now widespread illness that should be immediately controlled. At the end, a new designed mAb candidate is proposed based on our present *in silico* findings.

## THEORETICAL METHODS

Computational virology is an emergent research field that takes advantage of the progress from molecular and structural biology, immunology, bioinformatics and related areas to foster the understanding of virus, their evolutionary dynamics in nature, infectivity, pathogenesis, cell/host-tropism, viral assembly and their molecular interactions in general (including how to predict epitopes, how to design specific neutralizing antibodies and basically any drug design & discovery related to viral infections). ^27–34^ In particular, structural and interactive aspects can benefit from the solid foundations that computational molecular simulation methods such as Molecular Dynamics (MD)^35,36^ and Monte Carlo (MC)^35,37^ have achieved to probe the thermodynamic, dynamics and interactive properties of biomolecules in material science, food and pharma (see refs.^38,39^ for reviews). Here, we applied a fast constant-pH MC scheme^40,41^ for protein-protein studies^42,43^ to improve our understanding of the molecular interactions involving SARS-CoV-1 and 2 S RBD proteins and to identify key amino acids for the host-pathogen interactions.

### Molecular systems and their structural modeling

Several molecular systems were investigated in the present study employing the two SARS-CoV-1 and 2 S RBD proteins (see Figure 1) with ACE2 and the fragments of the mAbs 80R, CR3022, m396, and F26G29. Typically, these fragments of mAbs are fusion proteins from variable regions of the heavy and light chains of immunoglobulins connected with a short linker peptide. Additional calculations were carried out for the SARS-CoV-2 S RBD with a new proposed mAb based on CR3022. For most of these macromolecules, three dimensional crystallographic structures are available at the RCSB Protein Data Bank (PDB):^44^ a) the SARS-CoV-1 S RBD protein was extracted from the PDB id 2AJF (chain E, resolution 2.9Å, pH 7.5) where it was found complexed with ACE2 (chain A) − see Figure 2; b) the fragment of the Ab 80R was taken out from the PDB id 2GHW (chain B, resolution 2.3Å, pH 4.6); c) the anti-SARS-CoV-1 m396 Ab was extracted from the PDB id 2G75 (chains A and B, resolution 2.28Å, pH 8.5) removing part of the chains to keep only the variable regions and the short linker peptide; d) F26G19 Fab was taken out from PDB id 3BGF (chains L and C, resolution 3.0Å, pH 5.5), following the same procedure used for m396. Missing regions in these proteins were built up using the “UCSF Chimera 1.11.2” interface^45^ of the program “Modeller” with default parameters.^46^ Figure 3 shows their final three-dimensional structures as used in this work. All PDB files were edited before the calculations. Water molecules and hetero atoms were completely removed from all used files. The “UCSF Chimera 1.11.2” package^45^ was employed for all molecular visualizations and representations too.

**Figure 1.**
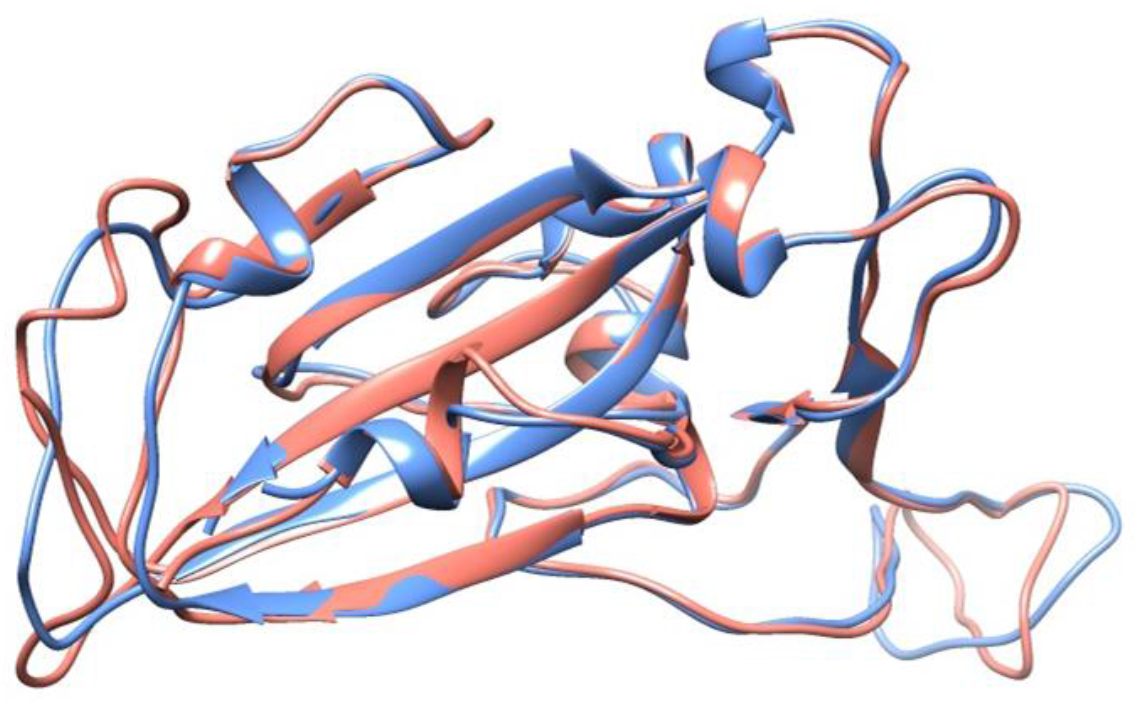
Crystal structure of the SARS-CoV-1 S RBD (PDB id 2AJF, chain E) and the modeled SARS-CoV-2 S RBD. See text for details regarding the modeling aspects. These macromolecules are shown, respectively, in blue and red in a ribbon representation. The RMSD between these structures is equal to 0.638Å.

**Figure 2.**
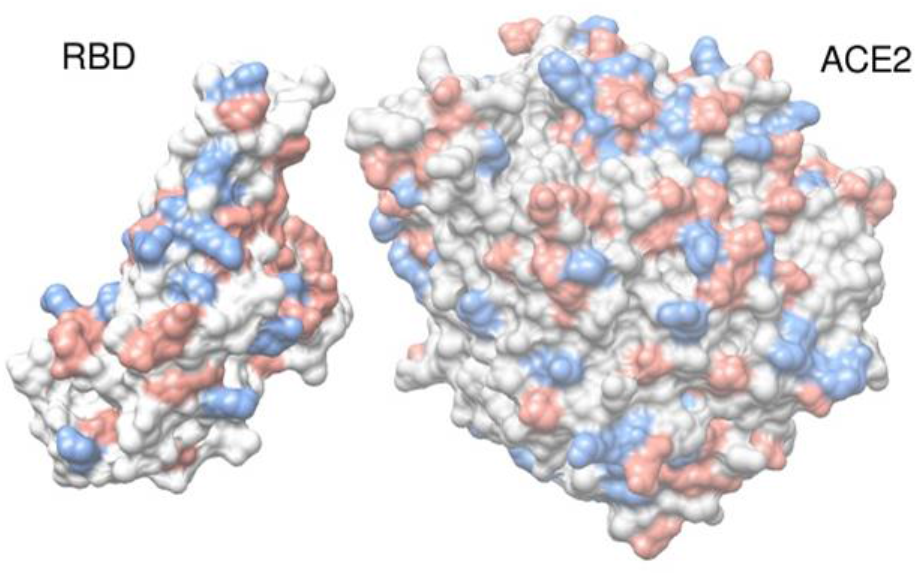
Crystal structure of SARS-CoV-1 S RBD complexed with ACE2 (PDB id 2AJF). Only standard amino acids of chain A (ACE2) and E (SARS-CoV-1 S RBD) are shown in a molecular representation using spheres for its atoms. Atoms are colored accordingly to their amino acids physical chemical properties: red for acid amino acids, blue for base amino acids and gray for non-titrating amino acids. For a better visualization of the interface, the ACE2 structure was translated ~12 Å.

**Figure 3.**
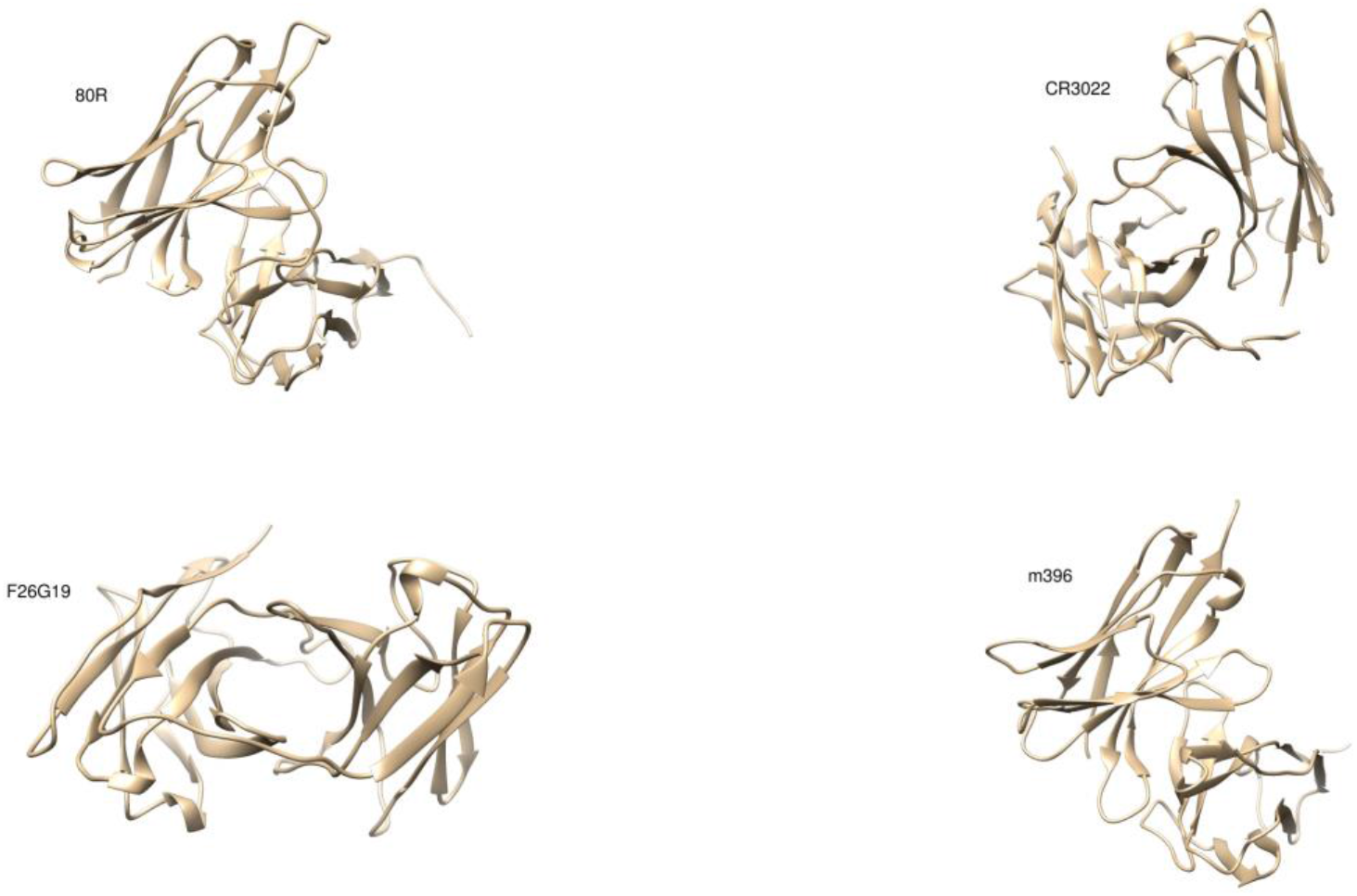
Molecular structures of the fragments of the investigated monoclonal antibodies. The fragments of the mAbs 80R (PDB id 2GHW), CR3022 (modeled – see the text for details), F26G19 (PDB id 3BGF) and m396 (PDB id 2G75) are shown in a ribbon representation.

When this study started, no experimental structure was available for the RBD of SARS-CoV-2 S protein. A model was built up at the SWISS-MODEL workspace (YP_009724390.1) based on the NCBI reference sequence NC_045512.^47^ The root-mean-square deviation (RMSD) of atomic positions between this modeled structure for the RBD of SARS-CoV-2 S protein and the available one for SARS-CoV-1 (PDB id 2AJF) is 0.638Å. The structural comparison between the RBD proteins of both SARS viruses can be seen in Figure 1. Recently, new experimental structures were solved. For example, a cryo-EM structure is now available for the prefusion S glycoprotein with a single incomplete RBD (PDB id 6VSB, resolution 3.46Å). The RMSD between our model and the S RBD (chain A) from this structure is 0.790Å. This number is closer to the RMSD differences between two chains of the same experimental (PDB id 6VSB) trimer structure (e.g. 0.668Å for chain A x chain B, and 0.732Å for chain A x chain C). Such diversity of possible conformations might motivate further studies exploring their effects on the theoretical predictions. These RMSD values also indicate that the modeled structure for the SARS-CoV-2 virus as used here is reasonable and within the expected conformational fluctuations from any other structure that could have been chosen for this work. Moreover, an intrinsic assumption here is that an experimental structure obtained at a given and specific physical chemical condition (ionic strength, pH, PEG6000 concentration, etc.) is valid in another condition.^27^

CR3022 is a particularly successful SARS-CoV-1 neutralizing human mAb first isolated from a convalescent patient by ter Meulen and co-authors.^48^ For the present study, its three dimensional structure (see Figure 3) was built up at the SWISS-MODEL workspace^47^ from the linear sequences of the variable regions of the heavy and light chains that were deposited in GenBank under accession numbers DQ168569 and DQ168570, respectively.^48^

### Molecular simulations

A large diversity of models is available for MD and MC molecular simulations.^38,49–52^ The need to repeat the calculations on several different physical chemical conditions and to obtain free energy of interactions at them drives the options to the so-called cost-effective coarse-grained (CG) models. These CG models offer the possibility to explore the main physical features of a system with a reduced number of parameters and lower computational costs.^27,42,43^ During the last years, a fast constant-pH (CpH) CG model has been devised to successfully study protein-protein interactions of several biological systems (including host-pathogens interactions).^27,42,43,53,54^ The possibility to fully consider the pH effects makes this modeling approach more appealing and appropriated to address this problem.^27,55^

A sketch of the simulations model is given in Figure 4. The S RBD proteins and the fragments of the mAbs were modeled as rigid bodies (i.e. bond lengths, angles, and dihedral angles are kept fixed) formed by a set of amino acids placed at positions given by their three-dimensional structures as described above. This additional approximation is justified by the prohibitive computational costs of constant-pH methods with pH-dependent conformational changes.^52,56,57^ Moreover, it is known that SARS-CoV-1 S protein does not exhibit large conformational changes upon the binding to ACE2 at least.^58^

**Figure 4.**
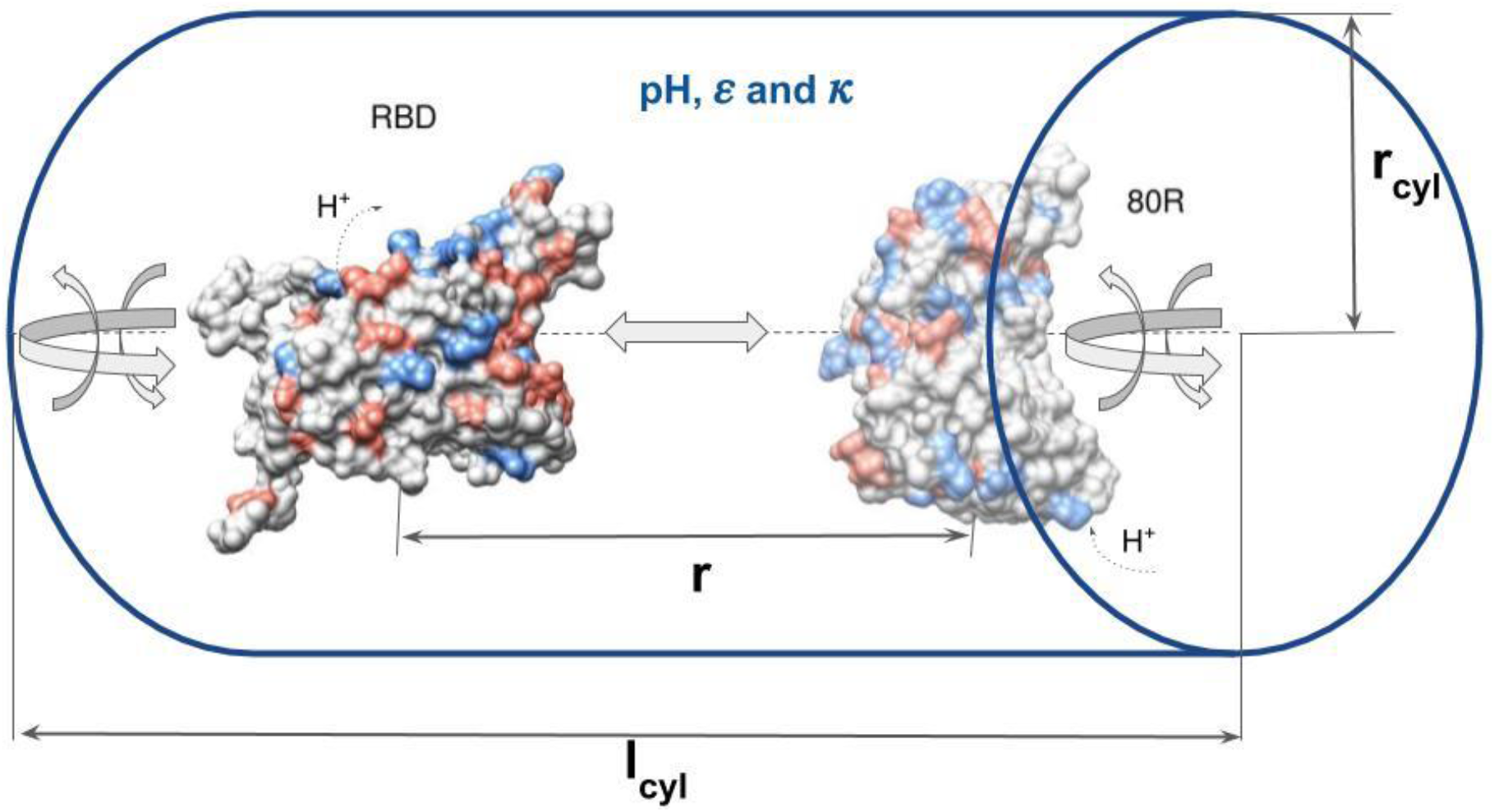
A sketch of the simulation model system for the constant-pH Monte Carlo simulations. A SARS-CoV-2 S RBD and the fragment of the mAb 80R (as given by the PDB id 2GHW) represented by a collection of charged Lennard-Jones spheres of radii *R*_*i*_ and valences *z*_*i*_ mimicking amino acids are surrounded by counter ions and added salt, implicitly described by the inverse Debye length *κ*. The solvent is represented by its static dielectric constant *ε*. Positive and negatively charged protein amino acids are represented in blue and red, respectively. The macromolecules’s centers of mass are separated by a distance *r*. The cylindrical simulation box is defined by the length *l*_*cyl*_ and radius *r*_*cyl*_. Translation (back and forward) and rotation (in all directions) possible movements are illustrated by the gray arrows while the protonation/deprotonation processes are indicated by the dashed arrows labeled with H^+^.

Each group of atoms that define an amino acid is converted in a single charged Lennard-Jones (LJ) sphere of radius (*R*_*i*_) and valence *z*_*i*_. This CG process turns a protein atomistic structure as a collection of charged LJ particles representing their amino acids. The centers-of-masses of the beads (mimicking amino acids) are used to place them accordingly to the coordinates given by the three-dimensional structures. The values of *R*_*i*_ for each type of amino acids were taken from ref.^53^. The valences of all ionizable residues are a function of the solution pH. The fast proton titration scheme (FPTS)^40,41,52^ was employed both to initially assign these valences *z*_*i*_′s for the amino acids and to let them vary during the simulation sampling at a given pH. This method has proved to predict p*K*_a_′s with a very good accuracy at low computational costs.^41^ The fundamental physical chemical basis of this titration scheme, its numerical implementation, benchmarks, discussions related to its approximations, pros and cons can be found in previous publications.^40,41,52,59^

As illustrated in Figure 4, two proteins are placed in an electroneutral open cylindrical simulation box, and free to translate back and forward along the axis in which their centers are laying, rotate in any direction and titrate. In this example, these two proteins are the modeled three dimensional structure of the SARS-CoV-2 S RBD and the crystallographic structure of the fragment of the mAb 80R. Unless otherwise specified, simulation runs were carried out with a cell of radius (*r*_*cyl*_) and height (*l*_*cyl*_) equals to 150 and 200Å, respectively. The static dielectric constant was set to 78.7 (assuming a temperature of 298K). Counter-ions and added salt particles were represented implicitly using a screening term, i.e., for two ionizable amino acids *i* and *j*, the screening is given by [exp(–*κ*r_ij_)] where *κ* is the modified inverse Debye length, and *r*_*ij*_ is the interparticle separation distance.^27,42,43,60^ Additionally, a simplified simulation box with only one protein present was used to characterize the titration properties of a single macromolecule.

The electrostatic interactions[*u*_*el*_(*r*_*ij*_)] between any two ionizable amino acids of valences *z*_*i*_ and *z*_*j*_are given by:

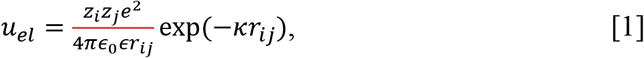

where *∊*_0_ is the dielectric constant of the vacuum (*∊*_0_ = 8.854 × 10^−12^C^2^/Nm^2^), *∊*_*s*_ is the dielectric constant of the medium (we used 78.7 to mimic an aqueous solution) and *e* = 1.602 × 10^−10^C is the elementary charge. See refs.^27,42,43,60^ for more details. Ionizable amino acids have their charged defined by the FPTS.^40,41^ All the others are fixed neutral.

Protein-protein interactions are also controlled by other physical contributions (van der Waals interactions, hydrophobic effect, and excluded volume repulsion).^27,42,43^ A simple and effective way to include their effects is by means of a LJ term [*u*_*vdw*_(*r*_*ij*_)] between the beads (amino acids).^27^ Mathematically, for any two beads (charged or neutral ones) *i* and *j*, *u*_*vdw*_(*r*_*ij*_) is given by

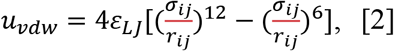

where *σ*_*ij*_ (= *R*_*i*_ + *R*_*j*_) is the separation distance of two amino acids *i* and *j* at contact. For instance, *σ*_*ij*_ for the pair VAL-GLU is 7.2Å (= *R*_*VAL*_ + *R*_*GLU*_, where *R*_*VAL*_ = 3.4Å and *R*_*GLU*_ = 3.8Å – see ref.^53^). The possibility to use different sizes for these beads allows the incorporation of non-specific contributions from the hydrophobic effect in the model.^43^ This should preserve the macromolecular hydrophobic moments^61^ and contributes to guide a correct docking orientation at short separation distances.^27^

The term *ε*_*LJ*_ regulates the strength of the attractive forces in the system.^27,42,43^ Typically, *ε*_*LJ*_ is assumed to be universal for any biomolecular system and equals to 0.124 kJ/mol.^42,43,53,62^ This should correspond to a Hamaker constant of ca. 9*k*_*B*_*T* (*k*_*B*_ = 1.380×10^−23^m^2^kgs^−2^K^−1^ is the Boltzmann constant, and *T* is the temperature in Kelvin) for amino acid pairs.^43,53,63^ However, this value might result in both an over or an underestimation of the attraction depending on the biomolecular system.^42,43,62^ For instance, *∊LJ* equals to 1.7*k*_*B*_*T* (a value 34 times greater than 0.124 kJ/mol) was necessary to reproduce experimental data for the histatin-5 adsorption to a hydrophilic silica surface.^62^ Conversely, the **β**-lactoglobulin–lactoferrin complexation seems to be overestimated by the usual value of *ε*_*LJ*_.^43^ Consequently, our research strategy has been to adopt the consensus value of 0.05*k*_*B*_*T* (= 0.124 kJ/mol) for *ε*_*LJ*_. This also implies that the outcomes should be interpreted with relative caution bearing in mind all the intrinsic approaches assumed in the modeling. The direct impact is seen in the free energy derivatives as discussed later at the results section.

Combining Eqs. (1) and (2), the total system’s interaction energy for a given configuration [*U*({*r*_*k*_})] can be written as:

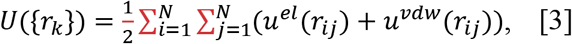

where {*r*_*k*_} are amino acid positions and *N* is their total number. This includes both charged and neutral beads.

This model was solved by Metropolis MC simulations that were performed at physiological ionic strength (150mM) and different pH conditions. The choice to simulate at pH 4.6 and 7.0 was motivated by the needs to understand the low and neutral pH conditions (e.g. low pH of endosomes). Furthermore, it seems controversial in the experimental works if the acidification is essential or not for uptake of cell-free SARS virus.^22,23,25^ The exact value of the acid pH condition is unknown. We made a choice to use the pH value of the crystallographic environment more acid among the studied structures (pH 4.6 for PDB id 2GHW). This also made possible to easily investigate the behavior of the systems at intermediate conditions by interpolation from the present outcomes.

After the proper equilibration of the simulated molecular systems, long production runs were carried out. Simulations whose focus was on titration properties [*Z*(pH) and p*K*_a_s] required 10^8^ MC steps. Conversely, runs to measure the free energy of interactions [*β*w(r)] were calculated from radial distribution functions [*β*w(r)=-ln g(r), where *β*=1/*k*_*B*_*T*] that demanded even longer runs with at least 3.0 10^9^ MC steps. These are massive simulations and very costly in terms of cpu time even at the CG representation. Four main factors contribute with this high cpu costs: *a)* the free energy barriers of the systems; *b)* the electrostatic coupling between a large number of titratable groups; *c)* the need to populate all the histogram bins used for the g(r) during the sampling; *d)* the reduction of the statistical noises in the calculated *β*w(r).^27,42,43^ Standard deviations were controlled by means of the use of 5 replicates per simulated system as done before for the study of flaviviruses.^27^

### Electrostatic Epitopes determined by the PROCEEEDpKa method

Antibody-antigen recognition is a challenger and intensive research field.^27,64–66^ It is a molecular process that involves different physical intermolecular interactions. Electrostatic interactions deserve a special attention in this process for several reasons [e.g. is long range nature, the fact that the interface antibody-antigen has a peculiar electrostatic pattern (richer in titratable groups) that is different than other general protein-protein interfaces,^65^ etc.].^27,55^ Such facts contribute to shift the canonical view of the “lock and key” (with a clear focus on the protein surface) to a broader definition that led to the “electrostatic epitopes” (EE) concept.^27^ This means that inner titratable groups (not only the ones at the epitope-paratope interface) can also participate in the interplay of interactions with Abs.

The EEs are the core idea of the PROCEEDpKa method^27^ where p*K*_a_ shifts are used to identify the key amino acids responsible for a host-pathogen association. It applies the fact that the location of these shifts is a practical mean to probe molecular interactions as before demonstrated.^67^ Moreover, this can be easily measured during computer simulations of a protein-protein complexation. The predictive properties of this powerful tool have been previously *i)* statistically analyzed for flaviviruses, *ii)* compared to other bioinformatic tools (that often ignore that pH and ionic strength can drastically affect the complexation process) and *iii)* discussed in details in a preceding work.^27^ The capacity of this method to test EE for *specific* mAbs makes it even more appealing for the present study where four known mAbs should be investigated. For the sake of convenience, predicted EE for the studied systems were graphically compared at the sequence level. The pairwise sequence alignments were generated by the server EMBOSS Needle^68^ with default settings.

## RESULTS AND DISCUSSION

### Free energy of interactions of SARS spike RBD proteins

One of the central questions in the understanding of the COVID-19, its pathology including the high transmissibility, and the possible therapeutic interventions to control the epidemics spreading is to decipher and prevent the molecular interactions between the S protein and ACE2.^25,69–73^ This SARS S protein-ACE2 complexation is the first step toward infecting the cell by the virus. Several studies have shown that both SARS-CoV-1 and SARS-CoV-2 viruses share the function interaction with this cell receptor (i.e. ACE2).^7,25,73,74^ We investigated the association pathway for the binding of the S RBD proteins to ACE2 for both SARS-CoV-1 and SARS-CoV-2 viruses by means of constant-pH MC simulations at two different solution pH values (4.6 and 7.0). The calculated free energy of interactions as given by the potentials of mean force [*βw(r)*] for these studied pH conditions at physiological ionic strength are given at Figure 5. Despite similar binding affinities observed in the present theoretical calculations (as seen in Figure 5) and in the laboratory experiments,^8^ the SARS-CoV-1 S RBD protein has a small tendency to bind to the ACE2 at both pH regimes. This agrees quite well with the experimental results measured by the biolayer interferometry binding (BLI) assay as reported by Walls and co-authors using the functional subunit of the S protein responsible for binding to the host cell receptor.^24^ The measured binding affinity (K_D_) was 5.0±0.1nM for the system SARS-CoV-1 S RDB(also referred to as the domain B^58^)–ACE2 and 1.2±0.1nM for the SARS-CoV-2 S RBD–ACE2. Yet, other experimental measurements using the S1 domain (this is the subunit that contains both the RBD and the N-terminal domain^58^) might suggest an inverted behavior where SARS-CoV-2 S1 domain would have a tendency for a stronger bind to ACE2 (K_D_=15.0±0.1nM)^75^ in comparison to SARS-CoV-1 S (K_D_=15.2nM)^8^. This small experimental difference of 0.2nM could be due to several reasons including the experimental uncertainties that were not reported in Tian’s work.^8^ In contrast to these results, another recent study^20^ advocated that SARS-CoV-2 has greater binding affinity for ACE2 than SARS-CoV-1. Even so, both the present theoretical and previously reported experimental data do agree that SARS-CoV-1 S RBD and SARS-CoV-2 S RBD have similar attraction to the ACE2. This high binding affinity implies that all human organs rich on ACE2 (oral and nasal mucosa, lung, stomach, small intestine, colon, skin, lymph nodes, thymus, bone marrow, spleen, liver, kidney, and brain)^76,77^ can be easily infected. A clear opportunity for the virus is the lung alveolar epithelial cells and enterocytes of the small intestine, where ACE2 is abundant.^76^

**Figure 5.**
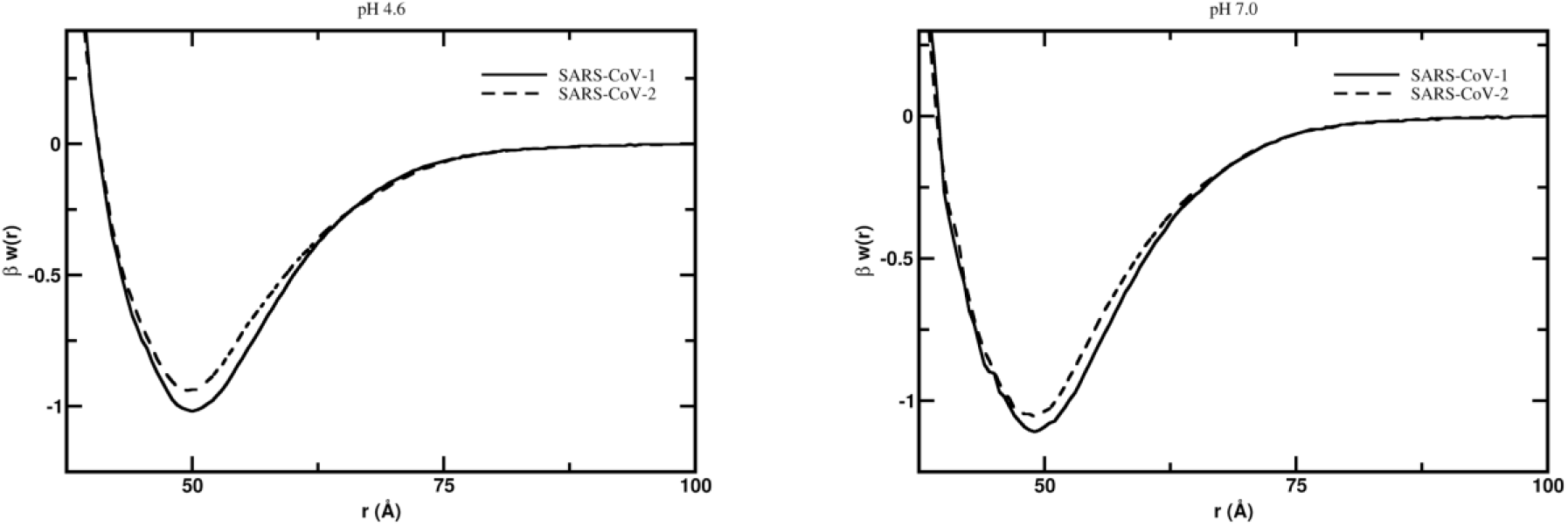
Free energy profiles for the interaction of RBD proteins with ACE2. The simulated free energy of interactions [*βw*(*r*)] between the centers of mass of the RBD proteins from both SARS-CoV-1 and SARS-CoV-2 and the ACE2 at different solution pH conditions. Salt concentration was fixed at 150mM. The structures of these macromolecules were extracted from the PDB id 2AJF for SARS-CoV-1 S RBD and ACE. SARS-CoV-2 S RBD was built-up by modeling as described in the text. Simulations started with the two molecules placed at random orientation and separation distance. Results for the systems SARS-CoV-1 and SARS-CoV-2 are show as continuous and dashed lines, respectively.

Note that the simulations were performed with a single RBD protein in the absence of the full structure of the S protein and the others two chains of the homotrimeric S glycoprotein. This brings with it the evidence that the other structural parts of the S1 subunit, the S2 subunit and the two other chains are not essential for the individual pair of RBD–ACE2 complexation. Also, this observation supports the argument that the dissociation of the S1 subunit complexed with ACE2 can happen without the interruption of the infection. This also allows the S2 subunit to transit from a metastable prefusion to its post-fusion state as a second step in the viral infection.^58,78^

At pH 4.6 which is closer to the low pH that occur outside the cell,^23,79^ the association between the SARS-CoV-1 S RBD protein and ACE2 showed a free energy depth [*βw*_*min*_] of −1.02 at the separation distance of 50.0Å (see Figure 1a). Conversely, for SARS-CoV-2 S RBD, *βw*_*min*_ is −0.95 at the separation distance of 49.5Å. The estimated standard deviations are 0.01*k*_*B*_*T* for all studied cases. Such computed measurements of *βw(r)* obtained by CG models that smooth the free-energy landscape must be interpreted with care as we already have pointed out above. By one side, it can be used a simple thermodynamic criterion that a negative free energy value (*βw*_*min*_<0) would result in a molecular complex. Conversely, any free energy value smaller than the thermal energy (1 *k*_*B*_*T*) would indicate an unstable association. The observed difference (*Δ*βw_min_) of 0.07 between the two RDB proteins is greater than the estimated statistical errors. However, we prefer to interpret such small differences as *tendencies* of the system. Based on previous studies^27,42,43,54,60^ where we have observed either that the computed complexation was weaker than the experimental measurements or too much stronger, we feel safer to use this data (and the others discussed below) in relative terms (i.e. comparing different situations). This allows us to successfully predict experimental observations respecting the limits of such CG models.^27,42,43,54^ The similarities between their free energy minima are relatively amplified at pH 4.6 (*Δ*βw_min_=0.07 as seen above) in comparison with pH 7.0 where *βw*_*min*_ is −1.11 and −1.06 (*Δ*βw_min_=0.05). Both viral S RBD proteins have their affinities to ACE2 slightly raised when pH is increased from the acid to the neutral regime. This somewhat higher binding affinity at neutral pH suggests that the role of pH for RBD proteins constantly under debate at the literature^22,25,73,80^ might not be so critical for the infection. It also reinforces the possibility that the viral cell invasion is *not* a pH-dependent process. Indeed, it seems that pH is more relevant for the next steps to continue the viral infection and not at the first entry level. This possible *non pH*-depend process might increase the opportunities for the SARS viruses to easily infect cells and therefore to contribute to its high infectivity. This might limit the use of chloroquine as an efficient drug against COVID-19 since its first action is to increase the cell pH. At least a neutral pH solution will not prevent the binding of the RBD to ACE2. On the opposite as it can favor this affinity.

Another important aspect is the evaluation of putative mAbs that could bind to the RBD of the new SARS-CoV-2. Following the work of Tian and collaborators,^8^ we investigated the interactions between the two SARS S RBD proteins with some of the most potent SARS-CoV-1 S RBD specific neutralizing antibodies (80R, F26G19, m396, CR3022). Figure 6 shows the free energy profiles at the acid regime and physiological salt conditions. For all studied fragments of Abs, a relatively stronger attraction is always observed for the S RBD protein from the SARS-CoV-1 interacting with any of these mAbs. This can be better seen in Figure 6b where the region around the well depth is highlighted. The lowest observed binding affinities are observed for the system SARS-CoV-2-F26G19 (blue dashed line, βw_min_=−0.63) followed by SARS-CoV-2-80R (black dashed line, βw_min_=−0.66) and SARS-CoV-2-m396 (green dashed line, βw_min_=−0.67). The difference between SARS-CoV-1-80R and SARS-CoV-2-m396 (*Δ*βw_min_=0.01) is within the estimated statistical errors. The most promising complexation was found for the SARS-CoV-2 RBD-CR3022 (red dashed line, βw_min_=−0.79) which is in good agreement with the experimental measurements.^8^ In fact, this was the only mAb that could bind potently with SARS-CoV-2 RBD (K_D_ of 6.3 nM determined by BLI assay) in the experiments performed by Tian and co-authors using the Ab isolated from the blood of a convalescent SARS patient.^8^ The Ab m396 only showed an insignificant binding at the highest measured concentration of 2μM in the experimental studies.^8^

**Figure 6.**
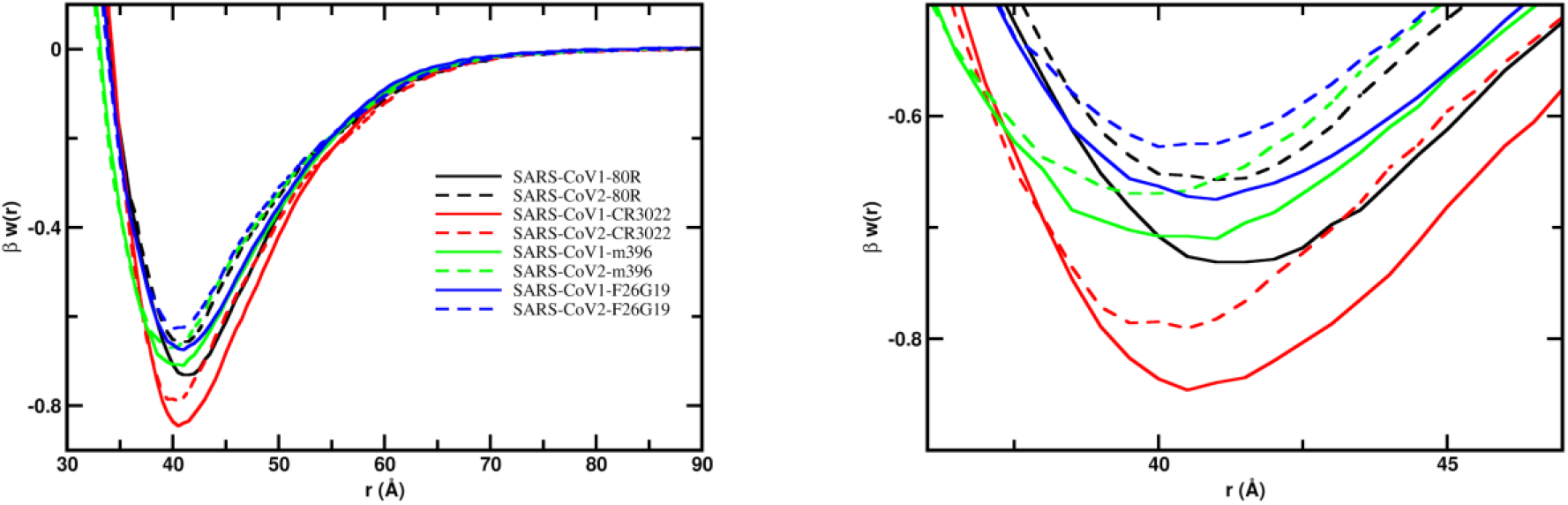
Free energy profiles for the interaction of RBD proteins with monoclonal antibodies. The simulated free energy of interactions [*βw*(*r*)] between the centers of mass of the RBD proteins from both SARS-CoV-1 and SARS-CoV-2 and the monoclonal antibodies at pH 4.6. Salt concentration was fixed at 150mM. See text for details about the structures of these macromolecules. Simulations started with the two molecules placed at random orientation and separation distance. Results for SARS-CoV-1 and SARS-CoV-2 are show as continuous and dashed lines, respectively. Different line colors are used for each fragment of the Abs: 80R (black), CR3022 (red), m396 (green) and F29G19 (blue). (a) *Left panel*: Full plot. (b) *Right panel*: The well depth region of the *βw*(*r*) for each studied complex.

Using the free energy minima values observed in the simulations, we can order the binding affinities for the RBD proteins from the lower to the higher as SARS-CoV-2-F26G19 (βw_min_=−0.63) < SARS-CoV-2-80R (βw_min_=−0.66) < SARS-CoV-2-m396 (βw_min_=−0.67) = SARS-CoV-1-F26G19 (βw_min_=−0.67) < SARS-CoV-1-m396 (βw_min_=−0.71) < SARS-CoV-1-80R (βw_min_=−0.73) < SARS-CoV-2-CR3022 (βw_min_=−0.79) < SARS-CoV-1-80R (βw_min_=−0.85). As mentioned above, the values of *βw*_*min*_ should be used in relative terms. Moreover, the work of Tian and co-authors suggested that only for CR3022 it was experimentally measured a reasonable binding.^8^ The combination of these information could indicate that a threshold of −0.67*K*_*B*_*T* for *βw*_*min*_ can be used to better refine the theoretical binding predictions of these macromolecular complexations (i.e. all viral protein-protein systems with a value of *βw*_*min*_ smaller than −0.67*K*_*B*_*T* are expected to experience binding *in vivo* at least). Table S1 summarizes the values of *βw*_*min*_ given between parenthesis.

It should be noted that the attraction between the S RBD proteins and ACE2 (*βw*_*min*_ equals to −1.02 and −0.95 for SARS-CoV-1 and SARS-CoV-2, respectively) is always stronger than what was calculated to any studied mAb including to the CR3022 (*βw*_*min*_ equals to −0.85 and −0.79 for SARS-CoV-1 and SARS-CoV-2, respectively) for both SARS viruses (see table S1). The same tendency was experimentally verified.^8^ It was measured by BLI assay a K_D_ of 6.3 nM for the binding of CR3022 to SARS-CoV-2 S RBD which corresponds to a fraction of 0.41 of the K_D_ measured for the binding of ACE2 to the same RBD (K_D_ equals to 15.2nM).^8^

### Physical chemistry properties

Next, we explored basic physical chemical aspects that could offer a simple and quick reasoning to understand the above free energy results and eventually be used as descriptors to scan databases of mAbs to filter promising ideal candidates. Although different driven forces can result in protein-protein complexation, ^42,43^ pH and charge-charge interactions seems especially important for viral proteins.^27,55,81–85^ Indeed, the protein net charges numbers (*Z*) obtained as function of the solution pH show that the SARS-CoV-2 S RDB protein is always slightly more positively charged than SARS-CoV-1 S RDB protein at the same physical chemical environment (*Z* equals to 5.2 and 5.5, respectively, for them at pH 4.6) − see table 1. Since all studied fragments of Abs are also positively charged at pH 4.6 (*Z* equals to 9.1, 4.2, 2.7, 5.8 for 80R, CR3022, m396 and F26G19, respectively), it can be easily seen that the order observed for the binding affinities above in the free energy analyses do follow a simple charge-charge rule for the mAbs with similar surface area (*A*~10,000 Å^2^). For the SARS-CoV-1 S RDB protein, from the weaker to the stronger repulsive cases in terms of the Coulomb contributions (Z_i_*Z_j_ assuming the same Bjerrum length, salt screening and separation distances^42,43,86^), the predicted order for the binding affinity is 80R (5.2*9.1=47.3) < F26G19 (5.2*5.8=30.2) < CR3022 (5.2*4.2=21.8). This agrees with the previous free energy analyses (see above). As large is *A*, larger is the attractive van der Waals interactions that can overcome the charge-charge repulsion. This can also explain why m396 (that is smaller and has roughly half of *A*) is less attracted to the RBD proteins even being slightly less positively charged (*Z* equals to 2.7 at pH 4.6) than the others (*Z*~5-6). Similarly, this is the physical reason to understand the stronger binding affinity that ACE2 (A=25,290Å^2^) has to the S RBD proteins. Although ACE2 (*Z*=5.4) and F26G19 (*Z*=5.8) have similar *Z*s, the molecular surface of ACE2 is ca. 2.5 times larger than F26G29 (A=10,120Å^2^). We tested the van der Waals (vdw) contribution comparing *βw*(*r*) for CR3022 and m396 in a model where all electrostatic interactions where completely switched off and only vdw interactions are considered. This test-case system is shown in Figure S1. It can be seen that m396 in this hypothetical test does have a weaker binding affinity to SARS-CoV-2 S RBD protein in comparison to CR3022 confirming the arguments above presented.

**Table 1.**
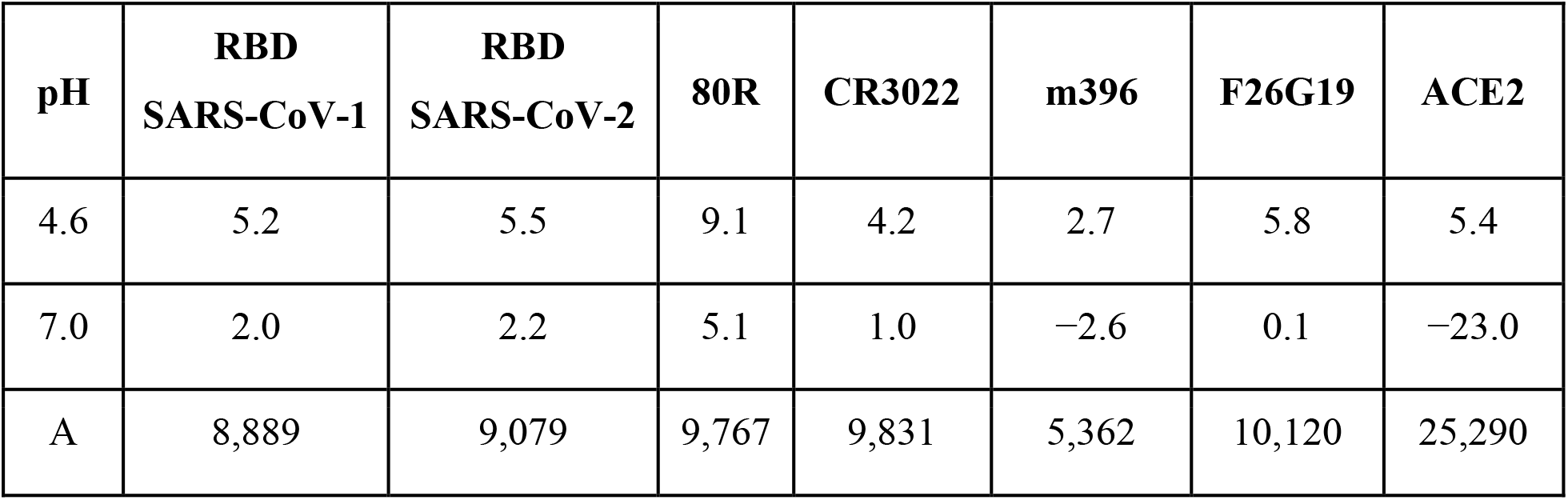
Main physical chemistry properties of the studied proteins. Protein net charge numbers (*Z*) for the investigated proteins at physiological ionic strength (150mM) and two different pH solution conditions (4.6 and 7.0). The macromolecular area (A) is given in Å^2^ as calculated by “UCSF Chimera” package.^45^

| pH | RBD<br>SARS-CoV-1 | RBD<br>SARS-CoV-2 | 80R | CR3022 | m396 | F26G19 | ACE2 |
| --- | --- | --- | --- | --- | --- | --- | --- |
| 4.6 | 5.2 | 5.5 | 9.1 | 4.2 | 2.7 | 5.8 | 5.4 |
| 7.0 | 2.0 | 2.2 | 5.1 | 1.0 | -2.6 | 0.1 | -23.0 |
| A | 8,889 | 9,079 | 9,767 | 9,831 | 5,362 | 10,120 | 25,290 |

### Insights to design a more efficient monoclonal antibody

Combining the findings above reported with a theoretical alanine scanning scheme employed to determine the contribution of specific titratable group to the complexation process, we identified three possible mutations that can improve the binding affinity of CR3022 to SARS-CoV-2 S RBD. The suggested mutations are K12E, K170A and R194A. These amino acids (K12, K170 and R194) can be seen in Figure 7 at the wild type structure of CR3022. The main physical chemical reasoning to design this new functional molecule was to reduce the net charge of CR3022 in general together with a decrease of the repulsion for groups that are closely located at the host-pathogen interface. Two amino acids substitutions (K170A and R194A) are suggested at this biological interface while the other one (K12E) is more peripheral (see Figure 7). Doing such mutations, the *Z* of the new molecule (labeled CR3022’) drops down from +4.2 to +1.2 at pH 4.6 and from +1.0 to −3.0 at pH 7.0.

**Figure 7.**
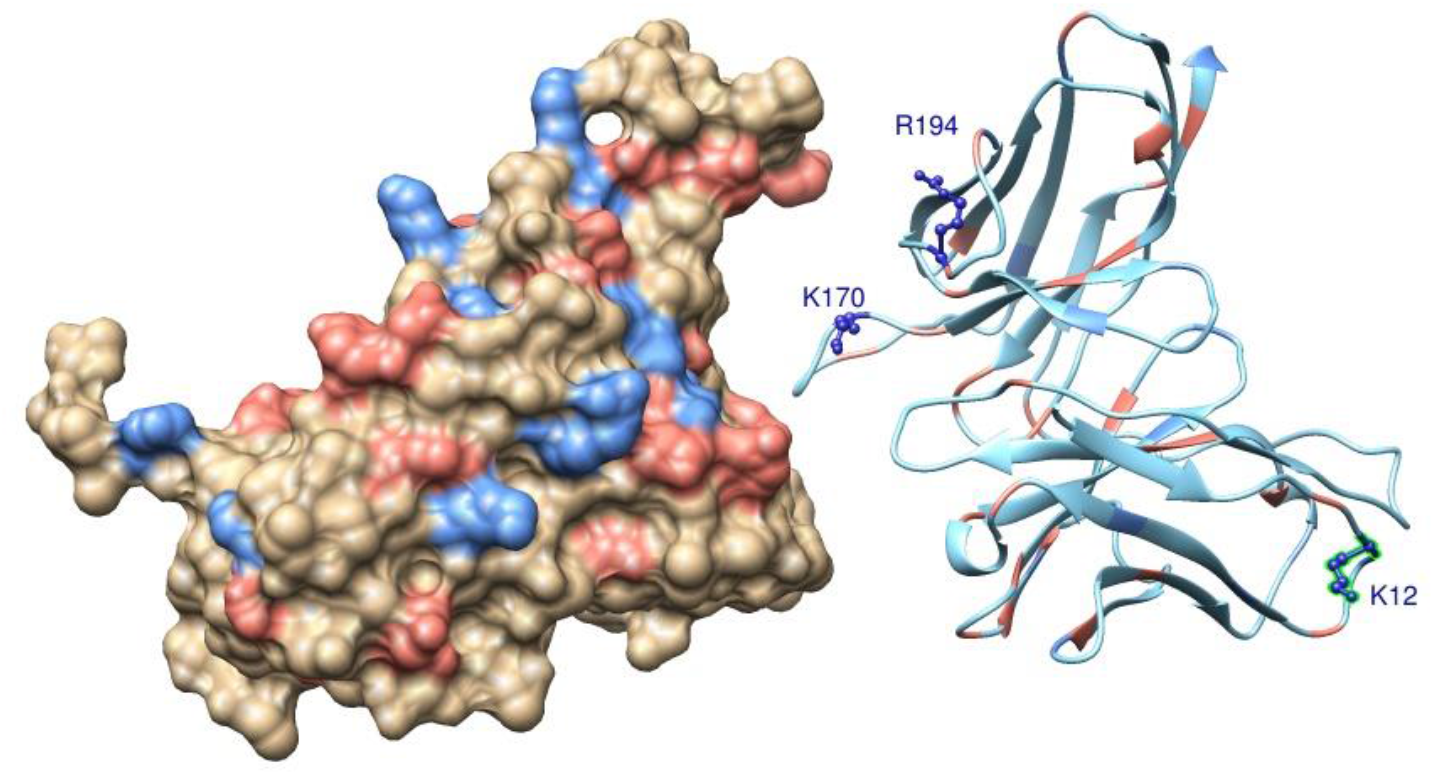
Molecular structures of a possible SARS-CoV-2 S RBD complexed with CR3022. Standard amino acids of SARS-CoV-2 S RBD (molecule at left) and CR3022 (molecule at right) are shown in a molecular representation using spheres and ribbons, respectively. Atoms are colored accordingly to their amino acids physical chemical properties: red for acid amino acids, blue for base amino acids and wheat/green for non-titrating amino acids. For a better visualization of the interface, the two macromolecules were placed ~12 Å apart from each other. Suggested residues to be mutated to improve the functional properties of CR3022 are indicated by the labeled amino acids (K12, K170, R194) are represented using the ball-and-stick model.

The binding affinity of this computer-designed molecule was tested. The calculated *βw*(*r*) for this new fragment of mAb is given in Figure 8. As it can be seen, CR3022’ is now able to bind with an equivalent binding affinity to what was observed for the SARS-CoV-1S RBD-CR3022 system. *βw*_*min*_ decreased from −0.79 to −0.85 recovering the value found for the SARS-CoV-1 S RBD-CR3022 case − see Figure 6 and Table S1. Therefore, this is a promising designed mAb candidate to be carefully and systematically examined in further experimental assays.

**Figure 8.**
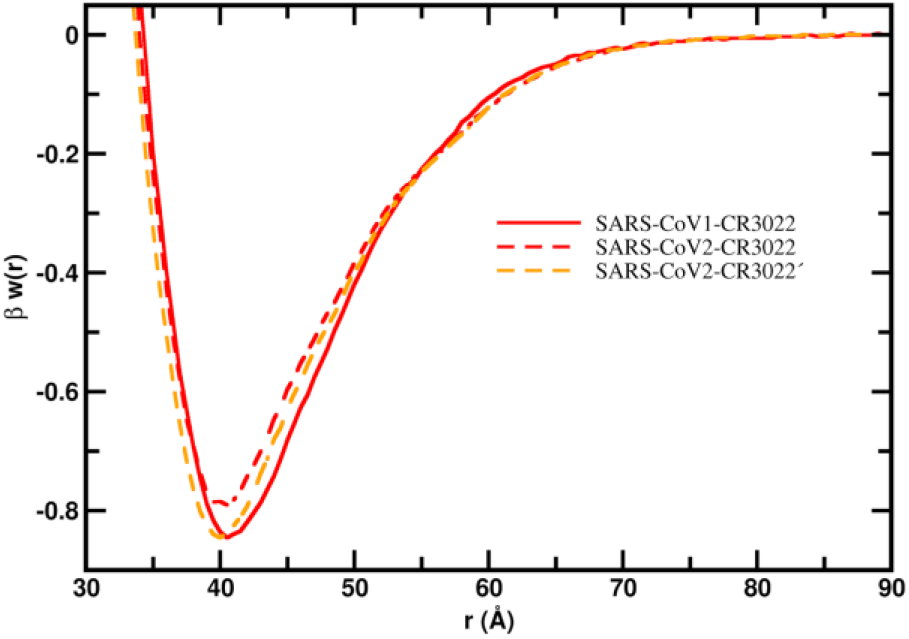
Free energy profile for the interaction of SARS-CoV-2 S RBD proteins with a new monoclonal antibody. The simulated free energy of interactions [*βw*(*r*)] between the centers of mass of the SARS-CoV S RBD protein and CR3022’ at pH 4.6 (dashed line in orange). Salt concentration was fixed at 150mM. See text for details about the structures of these macromolecules. Simulations started with the two molecules placed at random orientation and separation distance. The results for SARS-CoV-1-CR3022 and SARS-CoV-2-CR3022 (continuum and dashed lines in red) are also shown for comparison. This data was extracted from Figure 6.

### Estimates of the antigenic regions by p*Ka* shifts – The PROCEEDp*K*_*a*_ method

To refine this analysis at the sequence level, the PROCEEDp*K*a method^27^ was employed to determine the EE of the RBD proteins for the most relevant studied complexes. Three questions should be addressed here: 1) if SARS-CoV-1 and 2 S RBD proteins share a common binding region when they bind to ACE2; 2) if these viral RBD proteins interact with the mAbs using a similar epitope-paratope interface; 3) if the interaction with CR3022 and CR3022’ involves the same EE. The data to answer such questions is given in Figures 9 and 10.

**Figure 9.**
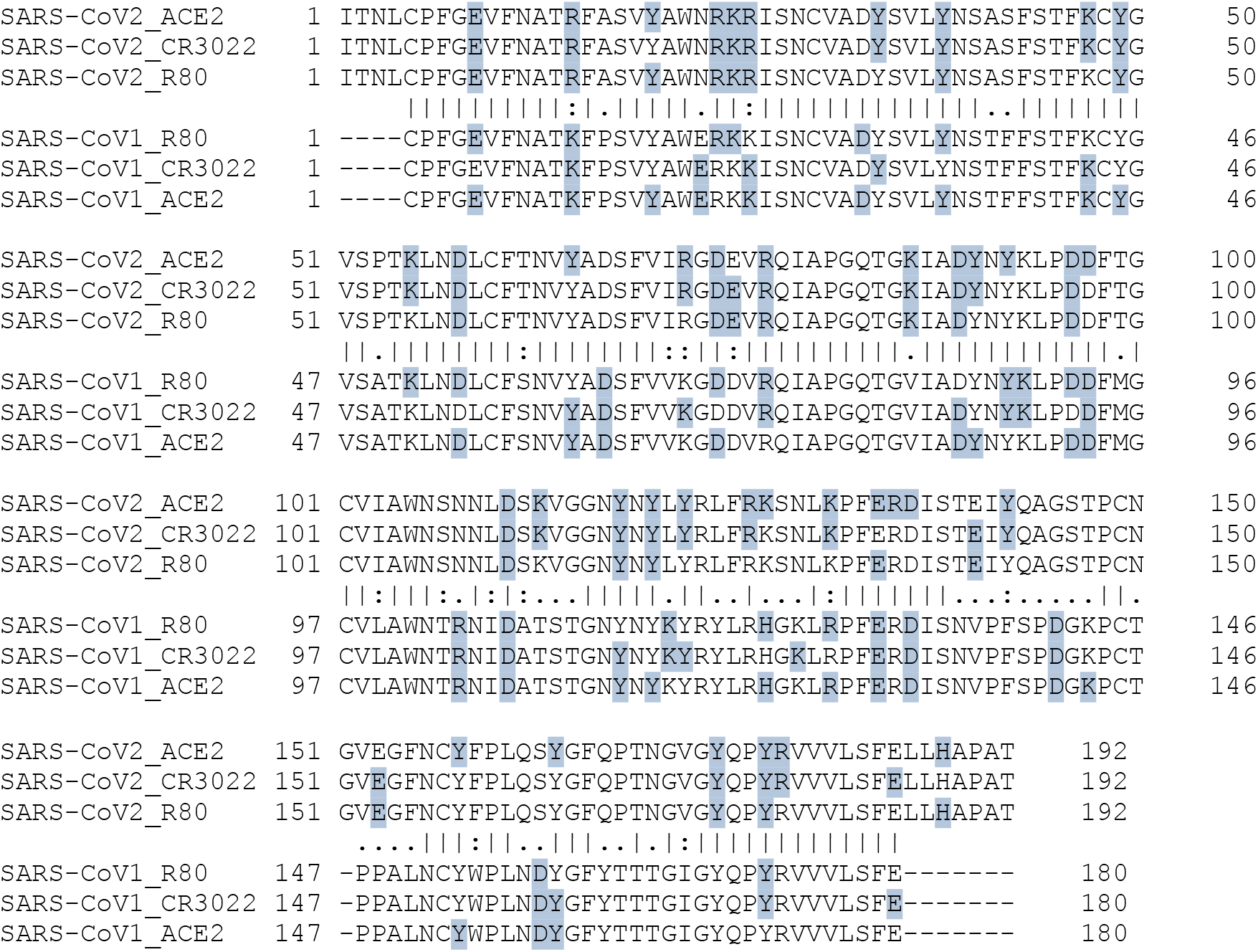
Electrostatic epitopes. Primary sequences of the SARS-CoV-1 S RBD and the SARS-CoV-2 S RBD with the interface with ACE2 and the estimated antigenic regions (shown in blue) for 80R and CR3022 by the electrostatic method. Data obtained using the threshold |Δp*Ka*|>0.01. Symbols between the two pairwise aligned sequences have the usual meaning: a) conservative amino acids where both sequences have the same residues are indicated by “|”; b) Similarities with a high score are marked with “;” and c) the ones with low positive score are indicated by “.”. Gaps are represented by “–”. Numbers are used to guide the identification of the amino acids sequence numbers.

**Figure 10.**
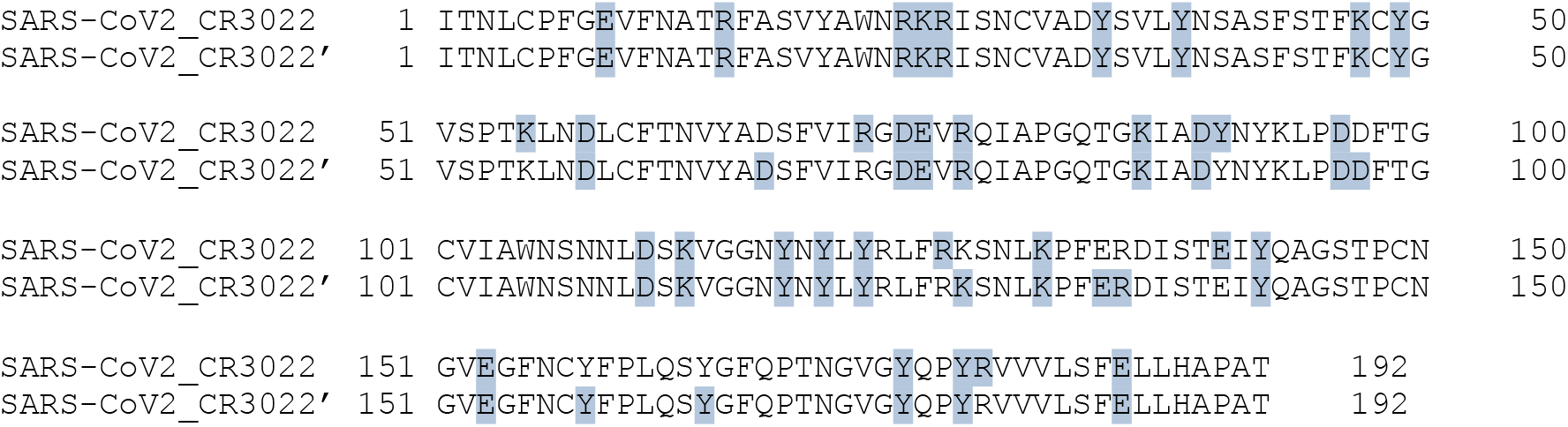
Electrostatic epitopes. Primary sequences of the SARS-CoV-1 S RBD and the SARS-CoV-2 S RBD with the estimated antigenic regions (shown in blue) for CR3022 and CR3022’ by the electrostatic method. Data for CR3022 is the same shown in Figure 9. All other details are also as in Figure 9.

In Figure 9, the primary sequences of SARS-CoV-1 and 2 S RBD proteins are plotted together with the estimated ionizable amino acids of the interface with ACE2 and the antigenic regions as defined by the PROCEEDp*K*a method. By the different distribution of amino acids identified as EE (shown in blue), it can be seen that the electrostatic method is sensitive to the structures and their titratable groups that can produce electrical perturbations on their partners when they are interacting as demonstrated before.^27^ The patterns observed for both viral SARS proteins are similar (i.e. they share a common region when they bind to ACE2) although some interesting observations can be made. Comparing the number of ionizable residues involved in the interactions for the RBD proteins of SARS-CoV-1 and ACE2 with the pair SARS-CoV-2 S RBD-ACE2, we can see an increase from 30 to 40 with a high number of common cases (22 amino acids) where the same amino acid interacts with ACE2 for both viral proteins. Most of the differences are observed for neighbor groups (e.g. “`AWERKKISN`” for SARS-CoV-1 and “`AWNRKRISN`” for SARS-CoV-2) indicating that the same biological interface was explored by the two viral RBD proteins in spite of their structural differences. The RBD protein responsible for COVID-19 clearly has more titratable residues interacting with ACE2 than its precursor. This observation *suggests* that its binding to ACE2 might be less specific than what happens for SARS-CoV-1. As such, the presence of an Ab may not completely block the SARS-CoV-2 S RBD-ACE2 interaction. In general, as seen in this Figure, most of the titratable groups from the viral RBD proteins involved in the binding to ACE2 are also the antigenic regions of the studied fragments of mAbs.

Virtually the same number of ionizable groups are seen at the antigenic regions for RBD proteins from SARS-CoV-1 (25 aa) and SARS-CoV-2 (24 aa) when interacting with 80R. The number of common cases is 12 while some regions are more affected by their structural differences (e.g. “DYSVLYNSTFFSTFKCYG” for SARS-CoV-1 and “DYSVLYNSASFSTFKCYG” for SARS-CoV-2). A replacement of an amino acid from the same physical chemical group (e.g. D by E) can be enough to result in different interactions (e.g. “`KGDDVRQIA`” for SARS-CoV-1 and “`RGDEVRQIA`” for SARS-CoV-2). CR3022 perturbed more titratable groups: 27 for SARS-CoV-1 and 33 for SARS-CoV-2. Taking into account what was hypothesized for ACE2 above, this might be an addition contribution to improve the capability of this mAb to interact and inhibit the RBD proteins. In fact, the experimental work of Tian and colleagues^8^ do show that the CR3022 binding to SARS-CoV-2 S RBD is not affected by ACE2. This might be the molecular basis for this behavior. We are careful with the use of stronger statements here due to the limitation of the theoretical approach. Several additional issues remain to be further investigated.

Finally, we compared the EE predictions for CR3022’ (34 aa) with CR3022 (33 aa) interacting with SARS-CoV-2 S RBD protein − see Figure 10. The predicted EEs for the interaction with CR3022’ are essentially the same ones observed for CR3022 (27 common aa). This implies that the suggested mutations here do not affected the antigenic regions. Another particularly interesting feature of this computer-designed molecule is that the number of EEs shared with ACE2 has increased from 18 (for CR3022) to 27 (for CR3022’). This might amplify the potential of this mAb candidate to better block the virus-host cell interaction.

## CONCLUSIONS

Free energies of interactions were calculated for several molecular complexes involving the RBD of SARS-CoV-1 and 2 spike proteins. The present theoretical results confirmed that both RBD proteins have similar binding affinity to the ACE2 as previously reported in experimental studies. This is observed at both acid and neutral pH regimes which probably indicates that the medium pH it is not so relevant for the beginning of the viral cell invasion. pH seems to be more important for the next steps of the viral infection and not at the first entry level. This has a direct implication for the drug development since the proposal of some like chloroquine is to raise cell pH.

Analyzing the interactions between these RBD proteins and the SARS-CoV-1 S RBD specific neutralizing mAbs (80R, F26G19, m396, CR3022) allowed us to reproduce the experimental results. The only mAb with measured affinities for the SARS-CoV-2 S RBD protein by BLI assay was CR3020^8^ which was also the one with higher affinity quantified in the present theoretical study. Moreover, we could map their electrostatic epitopes and identify that all mAbs tend to share the same titratable residues, and they are like the residues involved in the interaction with ACE2. However, the RBD protein responsible for COVID-19 clearly has more titratable residues interacting with ACE2 than its precursor suggesting that its binding to ACE2 might be less specific. This can explain the general difficulty that mAbs can experience to completely block the SARS-CoV-2 S RBD-ACE2 interaction.

Charge-charge interactions were found to be a good simple descriptor for a fast screening to the designing of improved mAb for diagnostics, therapeutics and vaccines. Our theoretical approach, while still being further developed, has identified three amino acids substitution that can increase the binding affinity of CR3022 to the RBD protein responsible for the present pandemic. These results can contribute to guide the design of new functional and high specific mAbs providing a cost-and-time-effective computational framework towards the development of better diagnostic strategies and an effective treatment and/or vaccine for COVID-19.

## ACKNOWLEDGMENTS

This work has been supported in part by the “Fundação de Amparo à Pesquisa do Estado de São Paulo” [Fapesp 2015/16116-3 (F.L.B.d.S.)] and the Conselho Nacional de Desenvolvimento Científico e Tecnológico (CNPq) [(F.L.B.d.S.)]. F.L.B.d.S. is also deeply thankful to resources provided by the Swedish National Infrastructure for Computing (SNIC) at NSC. It is also a pleasure to acknowledge initial discussions with Zhenlin Yang (Zhongshan Hospital, Fudan University, China) that also provided us some preliminary modelled structures used for our first tests. A. Laaksonen acknowledges Swedish Science Council for financial support, and partial support from a grant from Ministry of Research and Innovation of Romania (CNCS - UEFISCDI, project number PN-III-P4-ID-PCCF-2016-0050, within PNCDI III).

## SUPPLEMENTARY MATERIAL

Additional details are given as supplementary material for:

a. Values of the free energy depth for the interactions of the SARS-CoV-1 and 2 RBDs with different partners (ACE2, 80R, F26G19, m396, CR3022).
b. Van der Waals contributions to the free energy profiles for the interaction of RBD proteins with two monoclonal antibodies (CR3022 and m396).

**Table S1.** Values of the simulated free energy depth of the interactions for the association between the RBD proteins from both SARS-CoV-1 and SARS-CoV-2 and the monoclonal antibodies and ACE2 at physiological salt concentration (150mM). Data extracted from Figure 2. All values of βw_min_ are given in k_B_T units.

|  | **Macromolecular partner** |  |  |  |  |
| --- | --- | --- | --- | --- | --- |
| **RDBs** | **80R** | **CR3022** | **m396** | **F26G19** | **ACE2** |
| **SARS-CoV-1** | -0.73 | -0.85 | -0.71 | -0.67 | -1.02 |
| **SARS-CoV-2** | -0.66 | -0.79 | -0.67 | -0.63 | -0.95 |

**Figure S1.**
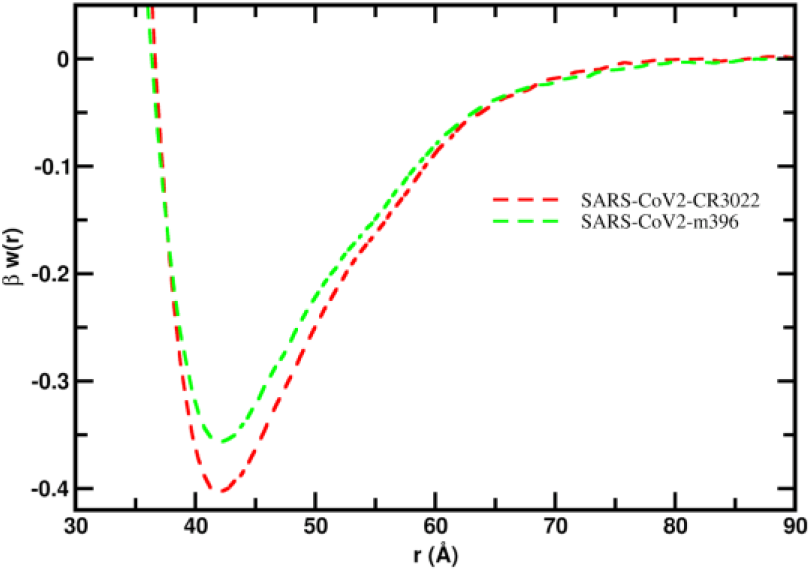
Van der Waals contributions to the free energy profiles for the interaction of RBD proteins with monoclonal antibodies. The simulated free energy of interactions [*βw*(*r*)] between the centers of mass of SARS-CoV-2 S RBD proteins and two monoclonal antibodies (CR30220 and m396) without the electrostatic contributions. All other details as in Figure 6a.

